## Supplemental files for "The *Chlamydia trachomatis* Inc Tri1 interacts with TRAF7 to displace native TRAF7 interacting partners"

```
CLUSTAL O(1.2.4) multiple sequence alignment

Tri1 (Serovar D)    MSFVGDSVPLRSYMPEAPLVDSASKARVSCCSERIAVLALGILSILFIVTGAALFIGAGW      60
Tri1 (Serovar L2)   MSFVGDSVPLRSYMPEAPLVDSASKARVSCCSERIAVLALGILSILFIVTGAALFIGAGW      60
                    ************************************************************

Tri1 (Serovar D)    TTLPMIDVVVTLVVFGSVMLGAVLTRISSYGGEPKKVSLDRFVLENERQGFLDKQRLADI      120
Tri1 (Serovar L2)   TTLPMINVVVTLVVFGSVMLGAVLTRISGYGGEPKKVSLDRFVLENERQGFLDKQRLADI      120
                    ******:**********************.******************************

Tri1 (Serovar D)    SKEEIALAKQQIEEEKEAILHSIFPND       147
Tri1 (Serovar L2)   SKEEIALAKQQIEEEKEAILHSIFPND       147
                    ***************************
```

**Fig S1.Alignment of Tri1 from L2 serovar D.**

A. Clustal alignment(61) of Tri1 from C. trachomatis serovars D and L2. Key: * , identical residues. :, highly conserved residues, ., weakly conserved residues.

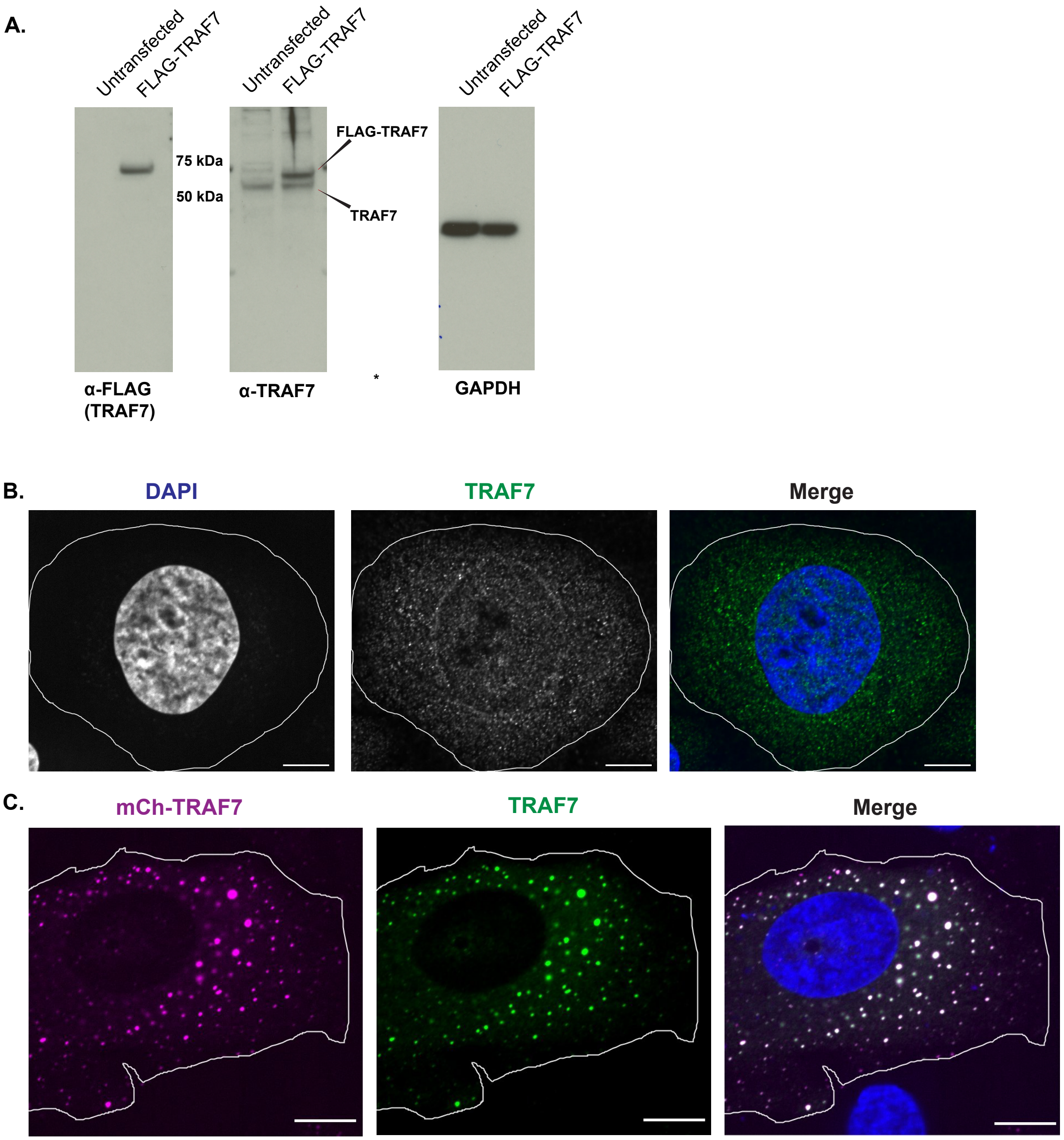


**Figure S2. TRAF7 antibody detects endogenous TRAF7 and ectopically expressed TRAF7.** A. Lysates prepared from a HeLa cell line that only expresses TRAF7 isoform 2 (see materials and methods) that were either untransfected or transfected with FLAG-TRAF7 were immunoblotted with the indicated antibodies. Antibodies to FLAG and TRAF7 detect similar sized proteins in the transfected lysates, which correspond with the predicted size of FLAG-TRAF7 (~78 kDa). The faster migrating band seen in the immunoblots probed with anti-TRAF7 represents endogenous TRAF7 isoform 2 (~67 kDa). GAPDH serves as a loading control. B. Hela cells that were transfected with mCh-TRAF7 for 24 hrs were fixed and stained with anti-TRAF7 and DAPI and imaged by fluorescence confocal microscopy. B. Endogenous TRAF7 localizes to both the nucleus and cytosol. Cell membrane is outlined. Shown are Single z-slices. Scale bar = 10 μm.

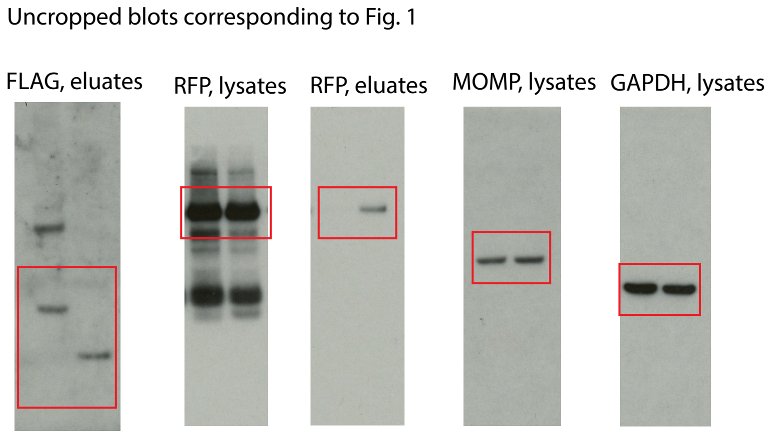


**Fig S3. Uncropped immunoblots corresponding to Figure 1.**

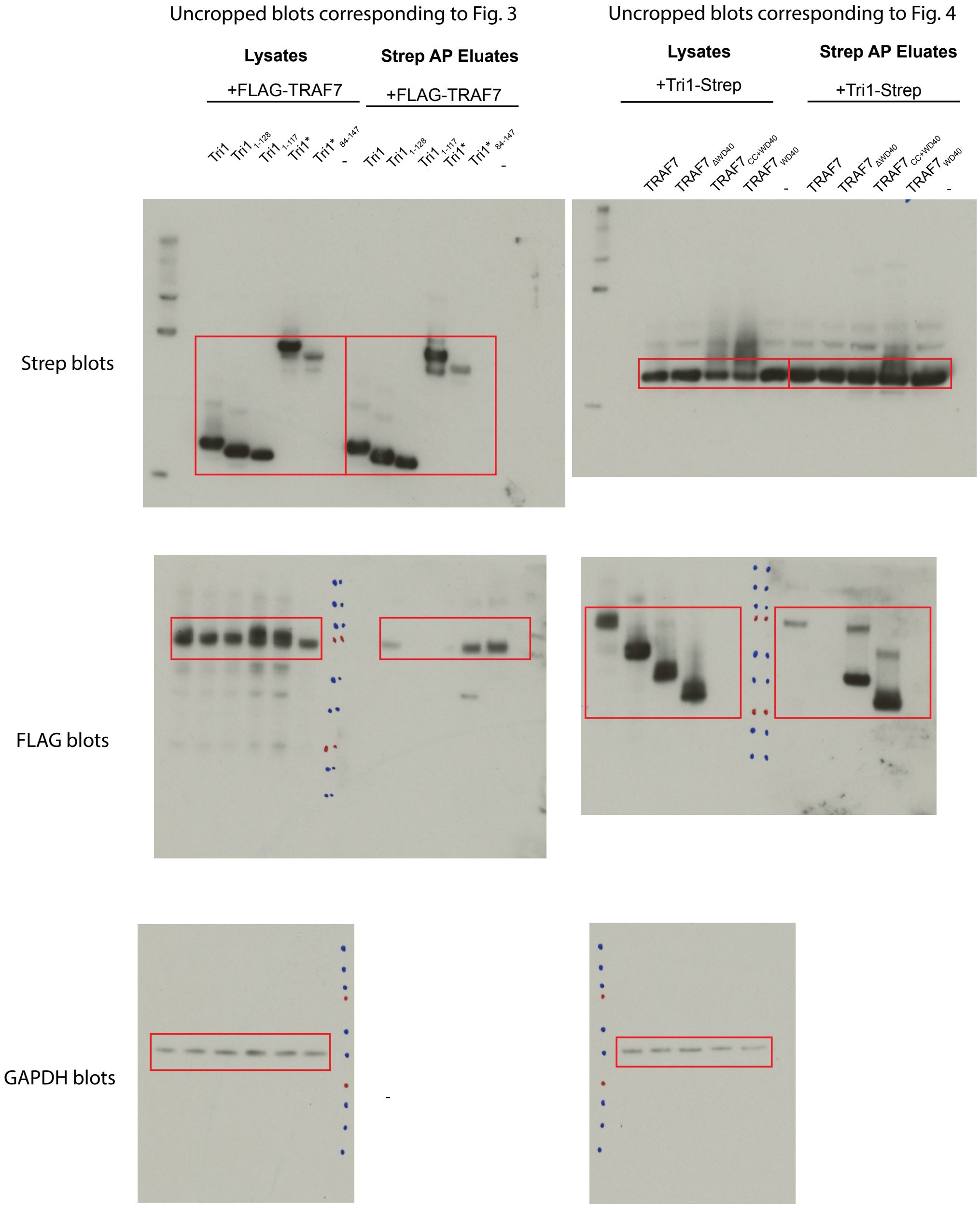


**Fig S4. Uncropped immunoblots corresponding to Figures 3 and 4.**

Uncropped blots corresponding to Fig. 5C

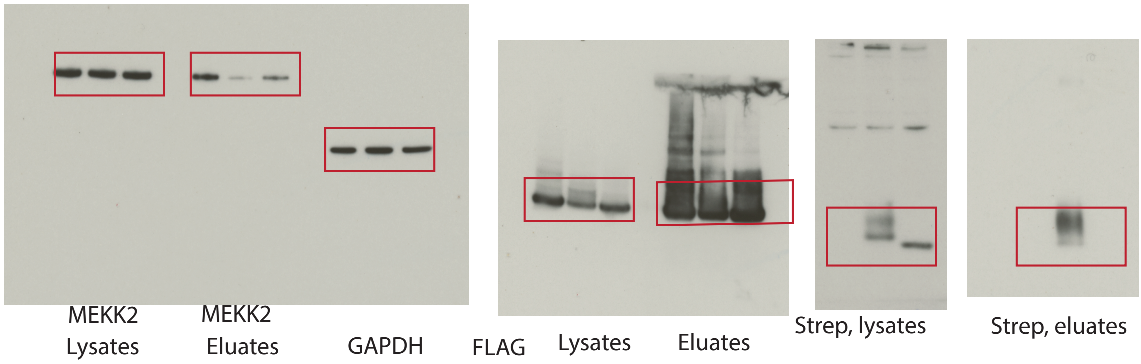


Uncropped blots corresponding to Fig. 5D

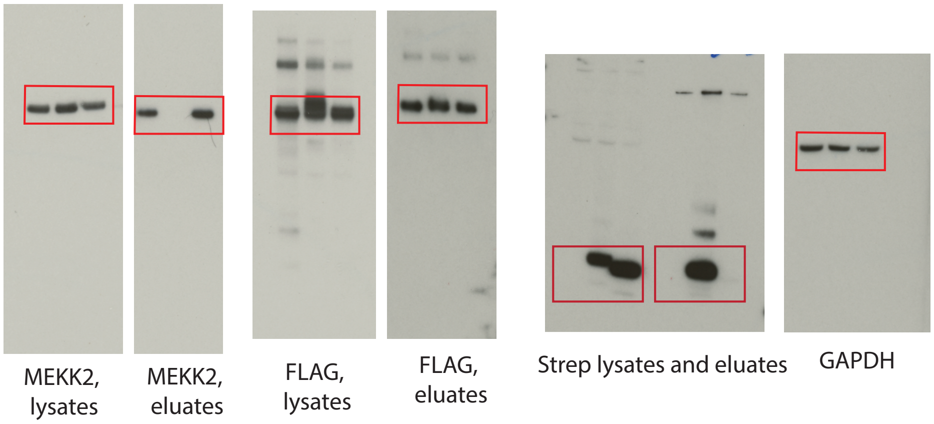


Uncropped blots corresponding to Fig. 5E

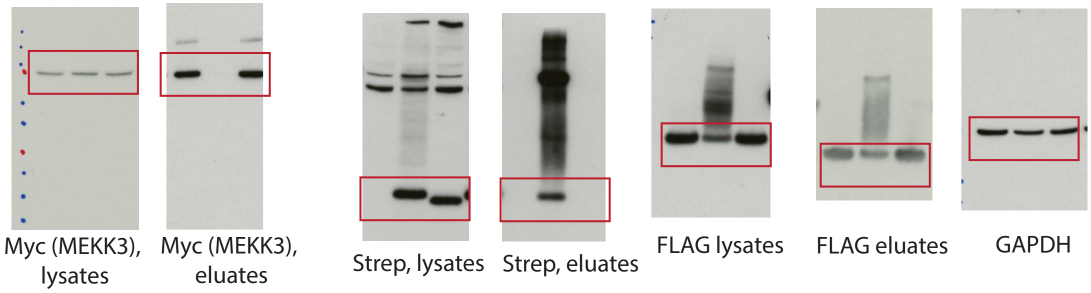


**Fig S5. Uncropped immunoblots corresponding to Figure 5.**
